# Mass Generation, Neuron Labelling and 3D Imaging of Minibrains

**DOI:** 10.1101/2020.07.11.198531

**Authors:** Subashika Govindan, Laura Batti, Samira F Osterop, Luc Stoppini, Adrien Roux

## Abstract

Minibrain is a spherical *in vitro* 3D brain organoid model, composed of a mixed population of neurons and glial cells, generated from human iPSC derived neural stem cells. Despite the advances in human brain organoid models, there is a lack of labelling and imaging methodologies to characterize these models. In this study, we present a step-by-step methodology to generate human minibrain nurseries and novel strategies to subsequently label projection neurons, perform immunohistochemistry and 3D imaging of the minibrains at large multiplexable scales. To visualize projection neurons, we adapt viral transduction and to visualize the organization of cell types we implement immunohistochemistry. To facilitate 3D imaging of minibrains, we present here pipelines and accessories for one step mounting and clearing suitable for confocal microscopy. The pipelines are specifically designed in such a way that the assays can be multiplexed with ease for large-scale screenings using minibrains. Using the pipeline, we present i. dendrite morphometric properties obtained from 3D neuron morphology reconstructions and ii. distribution and quantification of cell types in 3D across whole mount organoids.

## Introduction

*In vitro* culture models have been integral in studying aspects of brain development and function in real time. The advent of human induced pluripotent stem cells (iPSCs) has accelerated the development of miniature 3D human brain organoid models for the purpose of disease modelling, drug testing and molecular screening. Despite the advances in the field of brain organoid generation, the smaller size of organoids presents some challenges in performing classical histological and imaging techniques. On the other hand, some brain organoids are bigger in size but offer challenges for multiplexing for large scale screening studies and necrose relatively rapidly over a few months. Here, we present a protocol optimized for generating brain organoids termed as minibrains, where the size of the organoids and subsequent imaging techniques are optimized for large-scale screening studies at affordable cost, time and labor efficiency.

In this study, we present minibrains, a brain spheroid organoid model generated by non-directed differentiation of neural stem cells derived from human iPSCs (NSC^hiPS^) (Figure 1 and 2)^1^. By 6 weeks, minibrains display neuronal activity and by 12 weeks minibrains display synchronized neural networks (Figure 1)^1^. The size, simplicity and cost efficiency of generating minibrains make them an ideal choice for mass production for the purpose of large-scale screening and modelling studies (Figure 2, Supplementary Table 1). The average size of minibrains is around 550.64 (±) 75.19 μm and around 50 to 100 minibrains can be generated and maintained in one well of a 6 well plate (Figure 3, Table 1). The cost of generating approximately 100 minibrains is 33.50 CHF, the cost of generating and maintaining a “nursery” of approximately 6000 minibrains for about one-year costs about 3740 CHF. The smaller size of minibrains, allows efficient diffusion of nutrients to cells until the center of the organoids, reducing the occurrence of necrosis and maintaining organoid for up to 15 months and above. Minibrains can subsequently be maintained on an air-liquid interface (ALI) to facilitate brain on chip methodologies for neuronal network activity measurement using micro-electrode-array integrated bio-biochip (Supplementary Figure 1).

**Figure 1:**
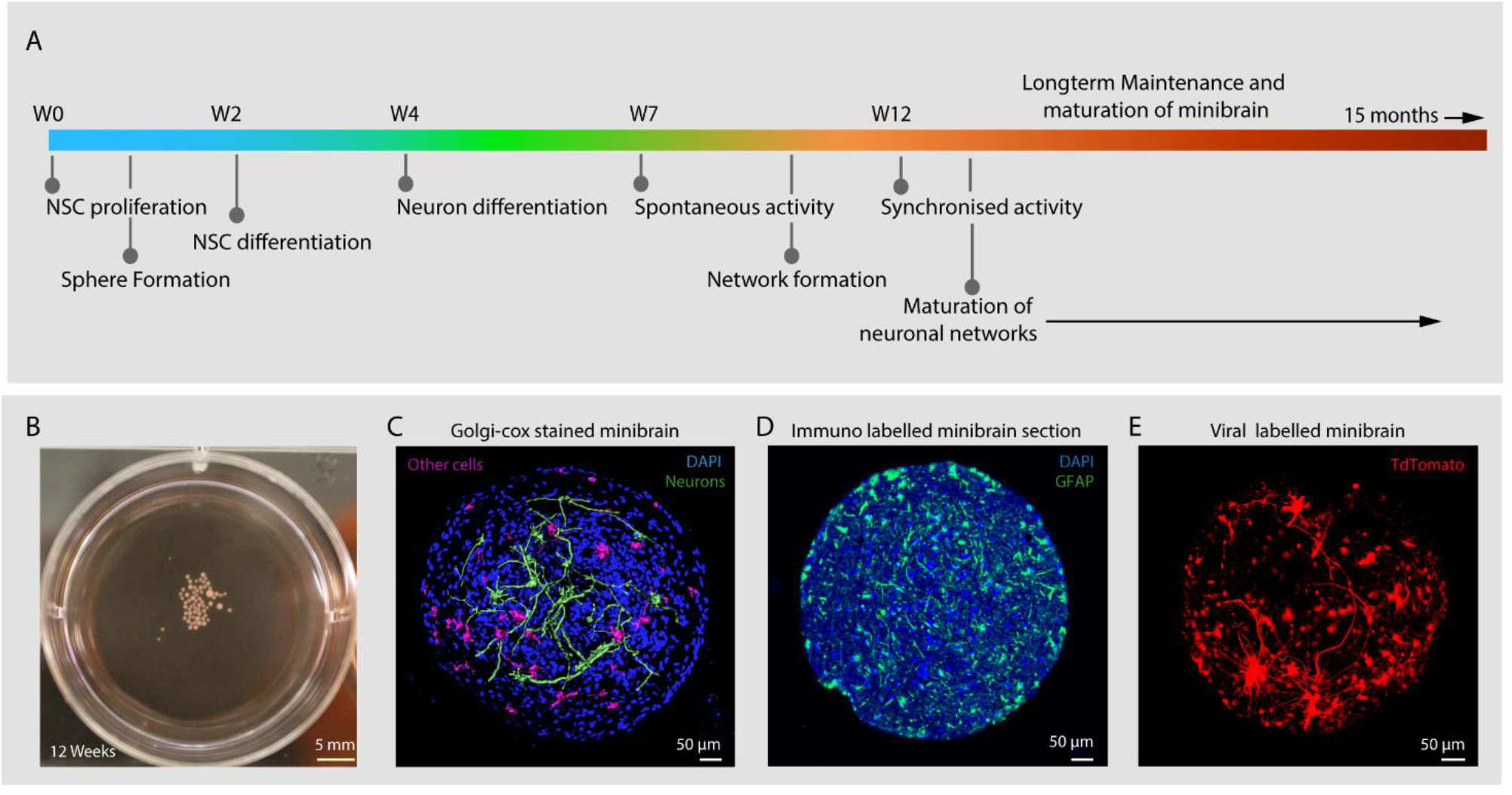
Introduction to minibrains. **(A)** Schematics show the timeline of processes involved in minibrain development. **(B)** shows 12 weeks old minibrains in a 6 well plate. **(C)** Volume rendered image of Golgi-Cox and DAPI stained whole minibrain showing neurons with elaborate projections (color coded in green), cells with short protrusions (color coded in pink), See supplementary video 1. **(D)** shows microtome cut minibrain section stained with GFAP, a marker of glial cells and DAPI (a nuclear stain). **(E)** shows volume rendered image of viral labelled whole minibrain showing tdTomato labelled projection neurons.

**Figure 2:**
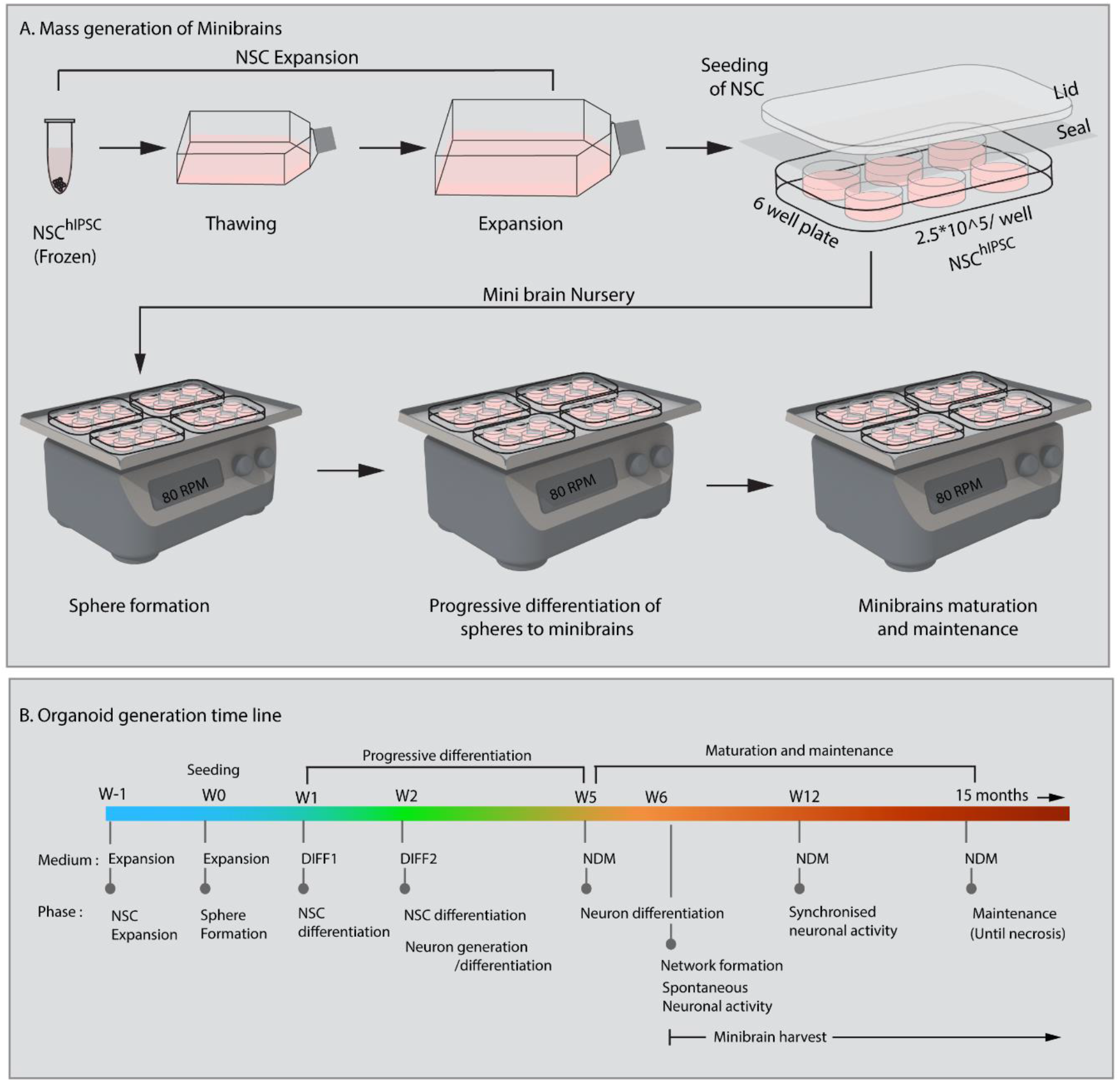
Mass generation of minibrains. **(A)** and **(B)** shows the processes and timeline of establishing and maintaining minibrain “nursery”.

**Figure 3:**
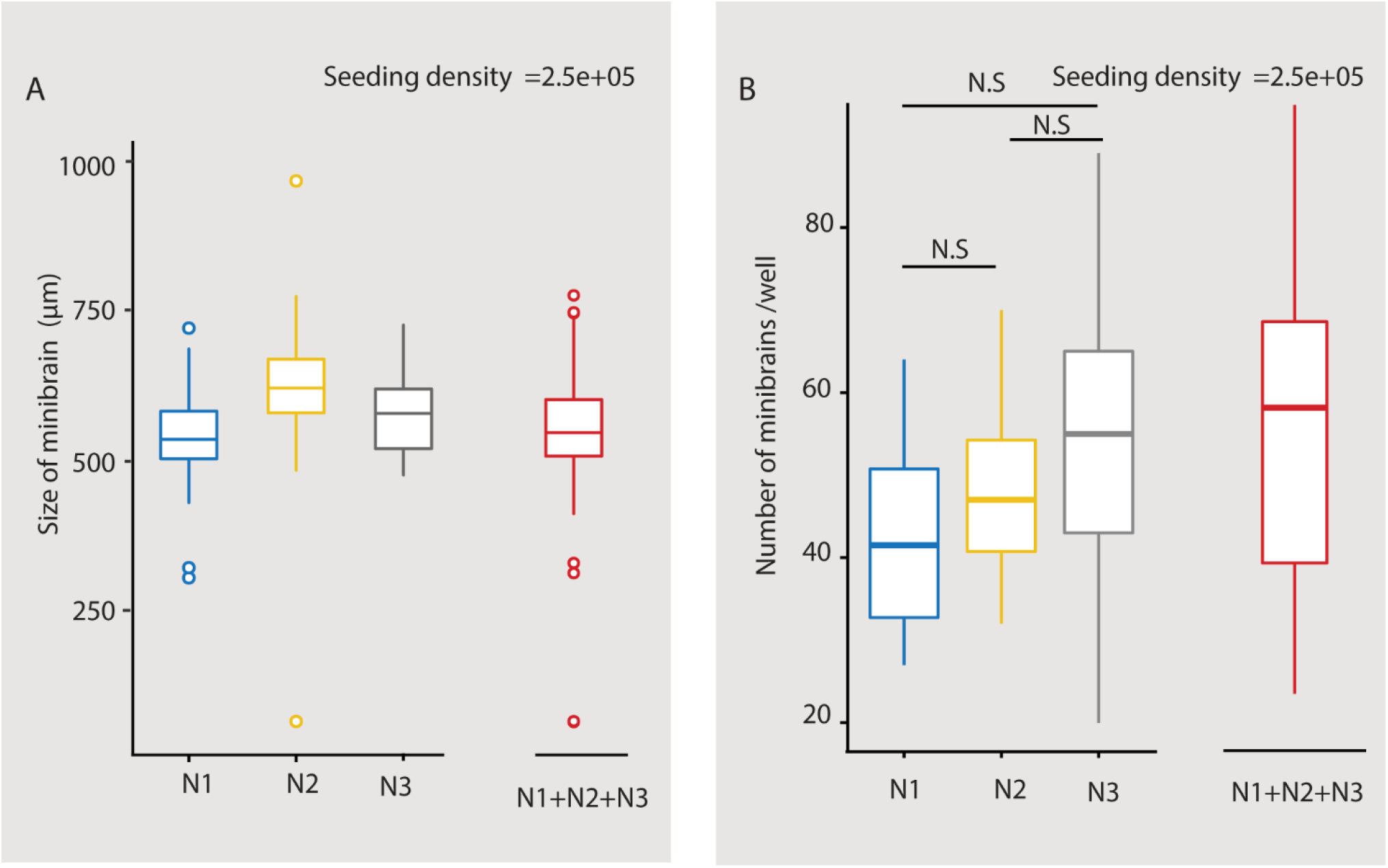
Size and number of minibrains generated using the protocol. **(A)** Shows the distribution of size of minibrains older than 8 weeks across three different batches (N1, N2 and N3) (with seeding density of 2.5e5 cells per well) as a whisker plot. Mean ± Standard deviation (S.D) of N1, N2 and N3 are 546.80 ± 62.40, 623.61 ± 68.52 and 584.73 ± 79.02 respectively. Collective mean ± S.D of all the three batches is 550.64 ± 75.19. **(B)** shows the number of minibrains generated per well in a 6 well plate across three different batches (with seeding density of 2.5e5 cells per well) as a whisker plot. The number of minibrains generated across three batches are not significantly different (p= 0.02 8) as per kruskal wallis statistical test. Mean ± S.D of N1, N2 and N3 are 42.93 ± 12.00, 47.75 ± 8.37, 52.47 ± 16.38 respectively. Collective mean ± S.D all the three batches is 48.97 ± 12.68.

**Table 1:** Plating guideline for Minibrain formation

|  | No. of cells | Medium volume | Average number of spheres | Volume of cell suspension |
| --- | --- | --- | --- | --- |
| 6 well | $1 \times 10^6$ | 3 ml | $95 \pm 34$ (N= 12 wells) | $2 \times 10^6 / X$ |
| 6 well | $2.5 \times 10^5$ | 3 ml | $49 \pm 13$ (N= 129 wells) | $5 \times 10^5 / X$ |
| 24 well | $2 \times 10^4$ | 500 μl | 1 or 2 | $4 \times 10^5 / X$ |

Morphological reconstruction of the neurons provides information about the dendrite morphometric and axonal properties that are crucial for network establishment. These properties can give valuable insight in large scale disease modelling and drug testing screens ^2,3^. Much is yet to be understood as to how human neuron morphology is driven in brain organoids where all connectivity pathways are miniaturized. Labelling of live neuron morphology involves artificial gene transfer techniques that enable expression of fluorescent reporters under the regulation of selective promoters. In previous studies, neurons were labelled in live brain organoids either through electroporation or viral infection of organoid slices, which are laborious processes ^4,5^. In our pipeline, we implement a novel strategy to label projection neurons with tdTomato fluorescence reporter in the minibrains using retrograde adeno associated viral particles (AAVrg) (Figure 6 B, C). Sparse neuronal labelling, together with a high signal to noise ratio of the labelled neurons, enabled 3D reconstructions of distinct neurons within the minibrain (Figure 6 B, C). For the first time, we show reconstruction of single neuron morphologies from a whole brain organoid, which would allow us in the future to understand neuron differentiation, diversity and connectivity in brain organoids (Figure 6 B, C, Supplementary figure 3).

3D imaging techniques are critical for reconstructing whole neuron morphology, assessing anatomical distribution of cell types and their interaction across brain organoids. Here we present a novel 3D imaging pipeline that allows imaging of whole brain organoids. 2D imaging methodologies require slicing of organoids which can be laborious, time consuming and leads to loss of tissue given the smaller sample size. In contrast, our 3D imaging pipeline relies on easily multiplexable one step non-invasive tissue clarification technique on whole organoids. We tested multiple tissue clarification protocols on our minibrains. While active CLARITY techniques are too harsh on the minibrains, passive CLARITY technique required embedding the organoids in agarose gel, a cumbersome process when processing multiple minibrains. Tissue clarification method using fructose-glycerol solution lead to distortion of the minibrain morphology (Figure 5 C). On the other hand, RapiClear™, a commercial tissue clearing agent, allowed direct mounting of minibrains without loss of morphology and best signal preservation for viral labelling, immunohistochemistry and Golgi-Cox staining (Figure 5–7, Supplementary video 1-4). We achieved efficient clarification by permeabilizing fixed minibrains using Triton X-100 before clearing with RapiClear™ (Figure 5 A, B and Section C and E). RapiClear™ allows preservation of mounted minibrains at 4°C and −20°C for long term storage. The cleared minibrains were compatible for both confocal and light sheet microscopy (Supplementary video 1-4, Figure 5). For high resolution imaging using upright microscopy, we have used an upright confocal imaging and designed sample holders and microscopy support to facilitate whole mount organoid imaging in high refractive index (RI) solution, while multiplexing up to 9 samples (Supplementary figure 2). The long travel distance of the immersion objectives (5.6 mm) allowed us to scan through the whole thickness of the cleared samples (Figure 5, Supplementary video 1, 3). The minibrains can be imaged until a depth of 150-250 μm using inverted microscopy allowing multiplexing up to 96 or 384 samples by using multi-well imaging plates (Supplementary video 5).

We present here an extensive step by step protocol for generation and maintenance of minibrain nurseries, ALI maintenance of minibrain, projection neuron labelling, optimized whole minibrain immunohistochemistry, one step mounting clarification, 3D imaging and design of imaging accessories that facilitate 3D imaging using confocal microscopy on cleared minibrains. We present dendrite morphometric properties, diverse reconstructed neuron morphologies, distribution of progenitors and POU3F2^+^ neurons in our minibrain (Figure 6 and 7, Supplementary Figure 4). Our pipeline is designed to facilitate the usage of minibrains as an *in vitro* 3D human neuronal model for large scale modelling and mass screening studies.

## Materials

### Reagents

GelTrex™ (ThermoFisher, #A1413301)

Stempro™ NSC SFM kit (ThermoFisher, #A1050901) containing

KnockOut™ DMEM/F-12 medium
StemPro™ Neural Supplement
FGF-basic (AA 10-155) Recombinant Human Protein
EGF Recombinant Human Protein

GlutaMAX™ Supplement (ThermoFisher, #35050038)

B27™ Plus Neuronal Culture System (ThermoFisher, #A3653401) containing

Neurobasal ™ Plus
B-27 Plus Supplement (50X)

Accutase (ThermoFisher, #00-4555-56)

StemPro™ hESC SFM (ThermoFisher, #A1000701) containing

DMEM/F12 + GlutaMAX™
Bovine serum albumin (BSA) 25%
Stempro^®^ hESC Supplement

Brain-Derived Neurotrophic Factor (BDNF) Recombinant Human Protein (ThermoFisher, #PHC7074)

Glial-Derived Neurotrophic Factor (GDNF) Recombinant Human Protein (ThermoFisher, #PHC7044)

Dibutyryl cyclic AMP (AMPc) (Merck Sigma, #D0627)

2-phospho-Ascorbic Acid (Merck Sigma, #49752)

RapiClear^®^ 1.47 (SUNJin Lab, #RC147001)

DAPI (4’,6-Diamidino-2-Phenylindole, Dihydrochloride) (Invitrogen, # D1306)

Triton X-100 (Merck Sigma Aldrich, # T8787-50ML)

Tween 20 (Merck Sigma Aldrich, # P9416-50ML)

1X Dulbecco’s PBS (DPBS) (ThermoFisher, #14040133)

Pierce™ 16% Formaldehyde (w/v), Methanol-free (ThermoFisher, #28906)

Trypan Blue Stain (0.4%) (ThermoFisher, #15250-061).

Purified Mouse Anti-Human Ki67 antibody (BD Bioscience, #556003)

Mouse Anti-Human POU3F2 antibody (DSHB, #PCRP-POU3F2-1A3-s)

Histodenz™ (Sigma, # D2158)

Goat anti-mouse TRITC (Abcam, #ab6786)

### Plastics and tools

6 well culture plate (Greiner Bio-One, #657 160)

96 well culture plate for imaging (Greiner bio-one, #655090)

384 well culture plate for imaging (Greiner bio-one, #781091)

24 well culture plate non-treated (Nunc, #144530)

Breathable plate sealer (Greiner Bio-One, #676051)

25 cm^2^ cell culture flask (T25) (Corning Falcon, #353109)

75 cm^2^ cell culture flask (T75) Flask (Corning Falcon, #353136)

175 cm^2^ cell culture flask (T175) Flask (Corning Falcon, #353112)

Sterile Hydrophilic PTFE membrane for tissue cultures, 2 mm diameter (named as « confetti ») (PTFE-005, HEPIA Biosciences)

Cell culture inserts for 6-well plate (Merck Millipore, #044003)

0.2 ml PCR tube (Thermoscientific, #AB-0784)

1.5 ml microfuge tubes (Eppendorf, #0030125150)

15 ml tube with conical bottom (Corning Falcon, #352096)

50 ml tube with conical bottom (Corning Falcon, #352070)

Pointe 10-20 μl cleanpack clearline sterile (Milian, #010320)

Pointe 20 μl cleanpack clearline sterile (Milian, #713178)

Pointe 200 μl cleanpack clearline sterile (Milian, #713179)

Pointe 1000 μl cleanpack clearline sterile (Milian, #713180)

10 μl pipette (Rainin, #17014388)

20 μl pipette (Rainin, #17014392)

200 μl pipette (Rainin, #17014391)

1000 μl pipette (Rainin, #17014382,)

2 ml sterile aspirating pipets (Corning Falcon, #357558)

12 mm glass coverslips (SPL Life Sciences, #20012)

Luna Cell Counting Slides (Logos biosystems, #L12001)

Sample holder (HEPIA, See supplementary material methods)

### Cells

NSC^hIPS^ (Human Neural Stem Cells derived from the human induced pluripotent stem (iPS) cell line) (ThermoFisher, #A3890101)

### Virus

AAVrg-CAG-tdTomato (codon diversified), titer value ≥ 7×10^12^ vg/ml (Addgene, #59462-AAVrg).

Important: Virus must be handled in biosafety level 2 (BSL2) facility

Note: Upon receipt of the virus, prepare 5 μl of aliquots in ice and store in −80°C.

### Equipment required

CO_2_ resistant Orbital Shakers (ThermoFisher, #88881102)

CO_2_ Incubator (ThermoFisher, #371)

LSM 880 confocal microscope (Zeiss)

20X/1.0 water immersion objective, with adjustable RI correction collar (RI 1.42 to 1.48) (Zeiss, #421459-9972-000)

TCS SPE-II microscope (Leica)

HC PL APO 10.0 × 0.30NA objective (Leica, #507902)

HCX PL FLUOTAR L 20X/0.40NA objective (Leica, #506242)

Specimen holder for cleared tissue imaging (HEPIA, See Supplementary materials and methods)

Laminar hood (SKAN AG, #MSF120)

Centrifuge (Eppendorf, #5804 R)

Media Warmer (Lab Armor™ Beads, #M706)

Elliptical shaker 3D Polymax 1040 complete (Heidolph Instruments, #543-42210-00)

Thermal cycler (MJ Research, #PTC-200)

Luna Automated Cell Counter (Logos biosystems, #L10001)

Inverted microscope (Zeiss, #Axiovert 25)

Laboratory vacuum pump (Milian, #886083)

Regine Horlogery watchmaker tweezers, type 7 (Beco Technic, #220337)

Mr. Frosty™ Freezing Container (ThermoFisher, #5100-0001)

### Reagent setup

#### GelTrex (1:200)

Thaw 1 ml vial GelTrex slowly at 4°C overnight. (!!! Attention: Agitation will cause clumping) Aseptically add 1 ml of GelTrex to 199ml of cold KnockOut ™ D-MEM/F12 medium. Can be stored at 4°C for up to 1 month.

### Stock solutions of supplements

Stock solution for EGF, FGF, BDNF (20 ng/ml), GDNF (20 ng/ml), and 2phospho-Ascorbic Acid (20 mM) are all prepared by resuspending the powder in DPBS 0.1% BSA, aliquot in 20 μl or 100 μl, freeze at −20°C and can be kept for 2 years.

Stock solution for Dibutyryl cyclic AMP (100 mM) is prepared by resuspending the powder in sterile water, aliquot in 20 μl or 100 μl, freeze at −20°C and can be kept for 2 years.

### Expansion Medium

Prepare expansion medium by aseptically mixing

500 ml, KnockOut ™ D-MEM/F12
5 ml, GlutaMAX ™ Supplement
10 ml, StemPro ™ Neural supplement

Before use, freshly and aseptically add 100 μl of EGF stock and 100 μl of FGF stock to 50ml of the above media mix.

### Differentiation 1 Medium (DIFF1)

For 50ml of medium mix aseptically

44.45 ml, D-MEM/F12 + GlutaMAX
3.6 ml, 25% BSA
1 ml, Stempro hESC Supplement
100 μl, BDNF stock solution
100 μl, GDNF stock solution
250 μl, Dibutyryl cyclic AMP stock solution
500 μl, 2phospho-Ascorbic Acid stock solution

Note: Do not overheat this medium BDNF and GDNF are sensitive to heat!

### Neuron Differentiation/Maintenance Medium (NDM)

1. Neurobasal ™ Plus 1x 500 ml
2. 50x B27 Plus Supplement 10 ml
3. GlutaMAX ™ Supplement 1.25 ml

### Differentiation 2 Medium (DIFF 2)

Mix DIFF1 medium and NDM medium at 1:1 ratio.

Note: Do not overheat this medium (for a long time) because several factors (BDNF-GDNF) are sensitive to heat!

### 4% PFA

Freshly dilute 0.5 ml of 16% PFA to 1.5 ml of 1X DPBS.

### Post fixation rinse buffer: DPBS-Tween

Dissolve 1 ml of Tween-20 in 100 ml of 1X DPBS. Store at 4°C for long time storage up until 6 months.

### Antigen retrieval buffer

Dissolve 1.5 g of sodium citrate (10 mM) in 450 ml water, adjust pH to 6 with 1 N HCl. Make up the final volume to 500 ml and add 480 μl of tween 20. Store at 4°C for long time storage up until 6 months.

### Blocking buffer

Dissolve 1 g of BSA and 2 ml of Triton X-100 in 100 ml of 1X DPBS. Store at 4°C for long time storage up until 6 months.

### Wash buffer

Dissolve 2 ml of Triton X-100 in 100 ml of 1X DPBS. Store at 4°C for long time storage up until 6 months.

### Primary antibody dilution

Dilute primary antibody in blocking buffer as per antibody manufacturer’s instructions.

Dilute Ki67 antibody (BD Bioscience, #556003) at 1:25 dilution in blocking buffer.

Dilute POU3F2 antibody (DSHB, #PCRP-POU3F2-1A3-s) at 1:5 dilution in blocking buffer

### Secondary antibody dilution

Dilute anti-mouse TRITC secondary antibody (Abcam, #ab6786) at 1:500 dilution in blocking buffer

### DAPI Stock solution

Make a 5 mg/mL DAPI stock solution by dissolving 10mg in 2ml deionized water. Make sure to dissolve the powder completely by vortexing the tube. Prepare aliquots and freeze at −20°C for long term storage. Protocol

### A. Generation of minibrains (Figure 2)

Before starting the experiment

1. Coat 1x T25 flask by adding 3 ml of GelTrex stock covering the surface area and incubate for 37°C for one hour. Note: optionally the coated flasks can be stored in 4°C for one month and pre warmed to room temperature before use.
2. Warm 9 ml expansion medium to room temperature for thawing and 5 ml at 37°C
3. Warm water bath at 37°C.

#### I. Thawing frozen aliquots of NSC^hIPS^

4. Thaw a vial of NSC^hIPS^ quickly in a 37°C water bath for not more than 30 seconds.
5. Transfer the contents of the vial to a 50 ml tube under the hood using a pipette.
6. Rinse the vial with 1ml prewarmed expansion medium (at room temperature) and add dropwise while swirling cells in a 50 ml tube.
7. Slowly add 8 ml of prewarmed expansion medium (at room temperature) while mixing gently.
8. Centrifuge thawed cells 250 g for 5 minutes at room temperature. A cell pellet will be visible after centrifugation.
9. Aspirate medium without disturbing the pellet and resuspend cell pellet in 5 ml expansion medium
10. Add the cell suspension in 5 ml of expansion medium to a GelTrex coated T25 flask and spread evenly
11. Every second day change the expansion medium.
12. When the cells reach 80% confluency, passage with Accutase (typically in 2-3 days) as described in step 13.

#### II. Passaging and expansion of NSC^hIPS^

Before starting the experiment: Warm Accutase and expansion medium (aliquots or the whole bottle) at 37°C. Coat GelTrex plates as described in step 1.

13. Aspirate all medium from flask and add 2 ml Accutase to flask and incubate at room temperature for 1-2 minutes.

Tip: Swirl the plate gently to better visualize the detachment of the cells.

14. To collect cells, add 2 ml of expansion medium to the detached cells and collect in a 15 ml tube.
15. Rinse the remaining cells with additional 2 ml of expansion and collect in the tube.
16. Centrifuge cells at 250 g for 5 minutes at room temperature.
17. Aspirate medium and resuspend pellet in 2-5 ml of expansion medium (depending on the size of the pellet).
18. Count live cells per ml on cell counter
  a. Mix 10 μl of Trypan Blue and 10 μl cell suspensions
  b. Transfer 10 μl of the mix on a counting slide
  c. Insert the counting slide in the Luna Automated Cell Counter and count.
  d. Number of live cells/ml = Number of counted cells* 2
19. Calculate volume of cell suspension needed for each T75 flask using the formula below.
  Number of cells required = 25*10^3^ cells/ cm^2^
  Surface area of T75 flask = 75 cm^2^
  Number of live cells/ml = X
  Volume of cell suspension required = Number of cells required * surface area of culture plate/ (X)
20. Add calculated volume of cell suspension and 10 ml of prewarmed expansion medium to GelTrex coated T75 flask.
21. Culture cells until they reach 80% confluency, typically in 2-3 days at 37°C 5% CO_2_.

#### III. Week 0: Formation of spheres (3D)

22. Prepare NSC^hIPS^ cell suspension and calculate the number of live cells per ml (X) as described in Step 13-18.
23. Calculate volume of cell suspension to be added based on the guidelines in Table 1 and as described in step 18.
24. Number of live cells calculated in step 22 is denoted as X in the table.
25. Add calculated volume of cell suspension and medium required based on the guidelines on table 1.
26. Seal the plate with breathable adhesive paper and close the lid.
27. To form spheres, place the cells on an orbital shaker at 80 rpm and culture for 7 days at 37°C 5% CO_2_. Note: spheres are apparent after 24 hours and increase in size is visible after 7 days due to proliferation. Important!!! From this point on the culture plates are always sealed with adhesive paper while culturing spheres in suspension.

#### IV. Week 1: Differentiation phase I

28. Position the plates at a slanted angle allowing spheres to settle down, aspirate the medium gently from the top without losing any spheres.
29. Add 2.5ml of DIFF1 medium per well (for a 6-well plate). Culture the spheres for 7 days at 80 rpm at 37°C 5% CO_2_.

#### V. Week 2-4: Differentiation phase II

30. Aspirate medium as described in step 27, Add 2.5 ml of DIFF2 medium (for a 6 well plate). Culture the spheres for 3 days at 80rpm in the cell culture incubator at 37°C 5% CO_2_.

#### VI. Week 3-5: Shifting to maintenance phase of the minibrains

31. Aspirate medium as described in step 27. Add 2.5 ml of NDM medium (for a 6 well plate). Change the NDM once per week and check the luminosity of the sphere (minibrain). **Troubleshooting:** Spheres can go through fusion upon shifting to maintenance medium (Figure 4). Shift one well of spheres to the maintenance medium, check for fusion of spheres after 24-48 hours. If spheres have fused in the test well, prolong differentiation phase II, repeat testing for fusion until spheres stop to fuse before shifting to maintenance medium. **Troubleshooting:** Health of the minibrains can be monitored by screening of necrosis in the minibrains (Figure 4). If a minibrain displays complete necrosis, discard the entire well of minibrains Minibrains can survive for 15 months and more, with neuronal activity. The spheres are termed as early minibrains starting from the 6th week, as that is when they display synchronized neuronal network activity. Mature activity is observed at 8 weeks.

**Figure 4:**
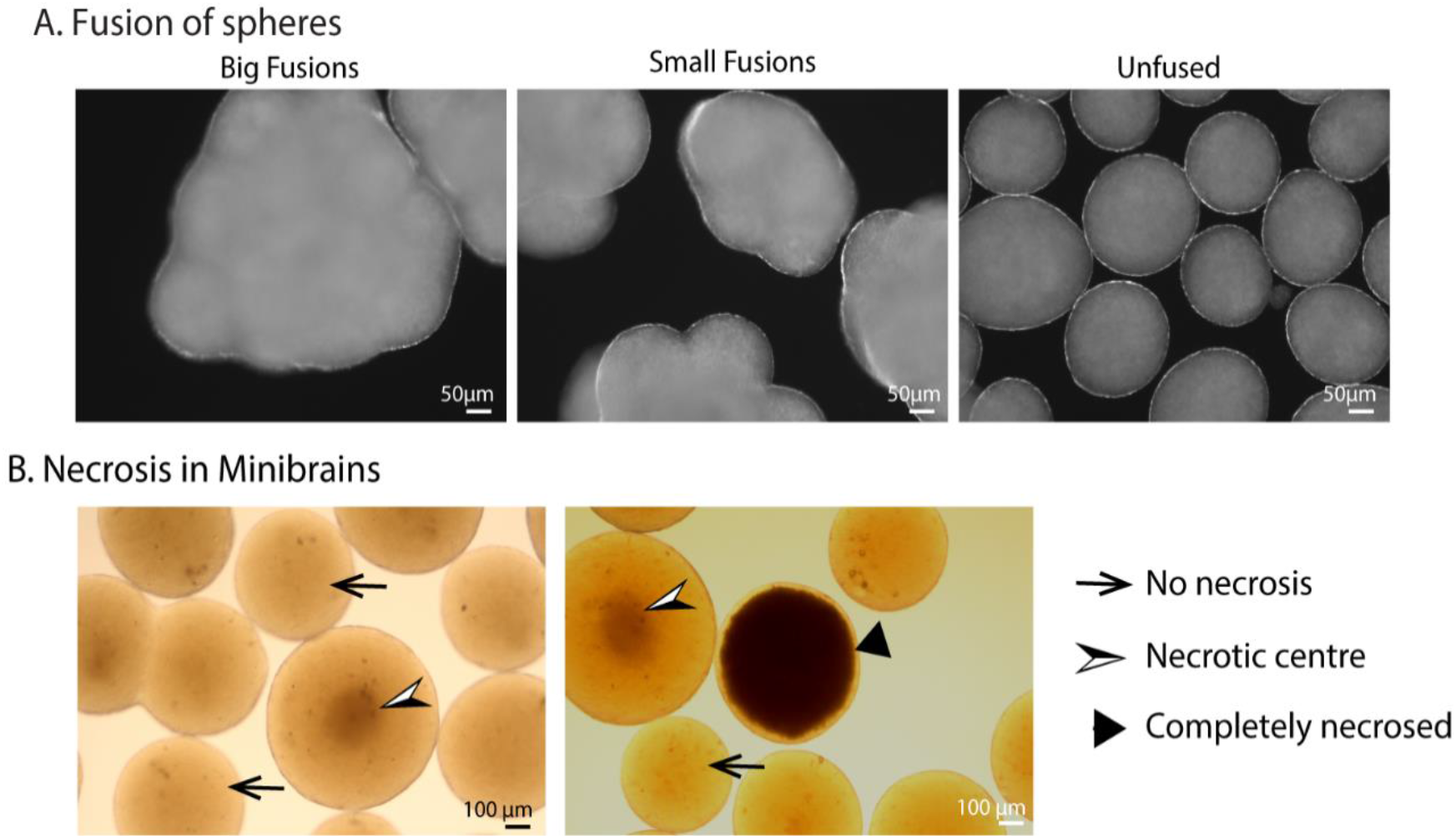
Fusions and necrosis during minibrain generation. **(A)** Left and Center images shows the extent of minibrain fusions. Image on the right shows unfused minibrains obtained after standardization. **(B)** shows monitoring of minibrain health by screening for necrosis.

**Figure 5:**
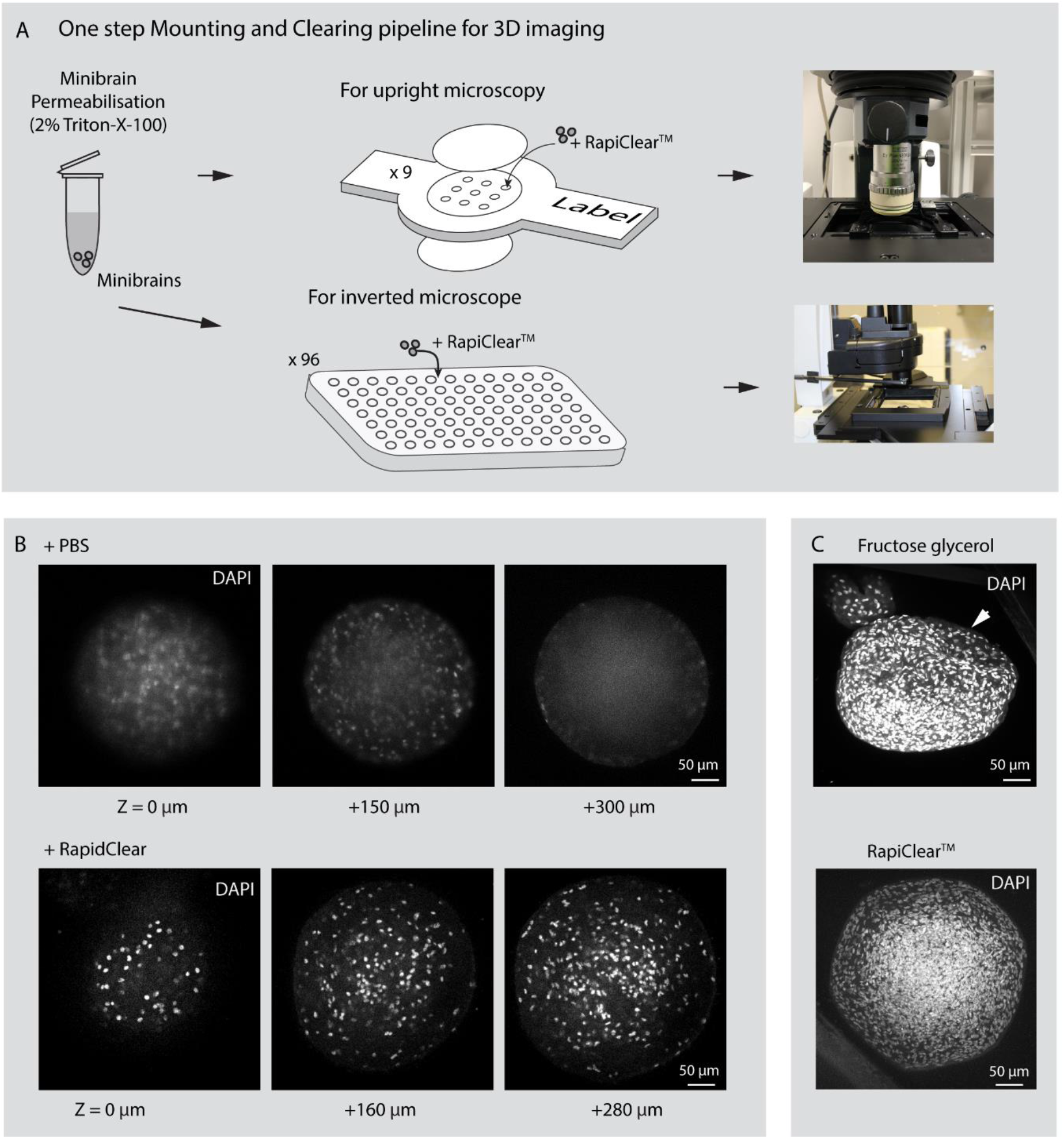
3D imaging of minibrains. **(A)** Shows the pipeline for imaging minibrains using upright and inverted confocal microscopes. **(B)** Shows the 2D optical sections of minibrains without clearing (top) and upon clearing with RapiClear™ at different Z stack/imaging depth. **(C)** Shows morphological changes in minibrains upon clearing with fructose glycerol (top) and intact morphology while using RapiClear™ (bottom).

#### VII. ALI culture of minibrain on confetti

32. Add 1 ml of NDM medium to a 6 well plate.
33. Place cell culture insert on the 6 well plate and place the confetti on the insert, enabling absorption of medium.
34. Warm the medium in the plate in the incubator for 30 minutes.
35. Set a 1000 μl pipette at 20 μl and transfer one sphere on the center of a confetti. (check the sphere for luminosity before transfer) Note: if the minibrains are big, cut the end of the pipette tip to transfer the sphere without damage.
36. Add 5 ml of water in the spaces between the well, to ensure humidity as the plates will not be sealed with breathable adhesive paper.
37. Change the medium twice per week by removing the maximum by aspiration and add 1 ml of Diff 3D medium.
38. Monitor necrosis under a light microscope.

### B. Viral labelling of projection neurons in live minibrains

Viral labelling must be done in biosafety level 2 equipped facility.

Before starting the experiment:

1. Prepare an ice box and place the viral aliquots in the ice to thaw
2. Set the centrifuge temperature to 4°C.
3. Warm 5ml of media in the cell culture incubator.
4. Spin the virus at 100 g for 10 minutes, to avoid loss of virus and any spill.

Viral infection of the minibrains (must be performed in a BSL2 facility under a sterile cell culture. Cut a 200 μl tip and transfer 3-4 minibrains to a 1.5 ml microcentrifuge tube. hood).

5. Remove the excess media
6. Add exactly 30 μl of media in each tube, let the minibrains settle to the bottom.
7. Add 0.5 μl of virus (3.5×10^9 particles) in each tube.

Note: The pipette tip must contact the minibrain while adding the virus. Do not mix the tube after the addition of the virus.

Using higher concentration of virus leads to more neurons labelled making it difficult to reconstruct single neuron morphologies.

8. Incubate the minibrains with virus in the cell culture incubator at 37°C 5% CO_2_ for 2-4 hours.
9. In the meantime, add 200 μl NDM medium to a 96 well plate and incubate at 37°C 5% CO_2_.
10. Cut the tip of a 200 μl pipette tip and transfer 1 minibrain per well in the prepared 96 well plate and culture the minibrains for a minimum of 48 hours.

### C. Immuno-histochemical staining of minibrains

#### I. Minibrain fixation

1. Remove media from the wells and add 100μl of 4% PFA for a 96 well plate, 1 ml of PFA for a 24 well plate and 3 ml of PFA for a 6 well plate and incubate for 45 minutes at room temperature.
2. Wash the minibrains with DPBS-Tween for 10 minutes 3 times.

#### II. Antigen retrieval

3. Transfer minibrains to PCR tube using a cut 200 μl tip. Remove excess DPBS-Tween, add 100 μl of antigen retrieval buffer and incubate at 95°C for 1 hour in a PCR machine.
4. Wash minibrains with minibrain wash buffer for 5 minutes.

#### III. Permeabilization and blocking of the minibrain

5. Using a cut 200 μl tip transfer minibrains to a 96 well plate and remove excess buffer.
6. Resuspend the minibrains in 70 μl of blocking buffer and incubate for minimum 4 hours at room temperature or overnight at 4°C on a rocker at 80 rpm

#### IV. Primary and secondary antibody labelling

7. Remove the blocking buffer and add 70 μl of primary antibody dilution. Incubate the minibrains in primary antibody for 4°C on a rocker at 80 rpm for 48 hours to 72 hours.
8. Remove the primary antibody, add 70 μl of wash buffer, incubate for 30 minutes on a rocker at 80 rpm. Repeat two more times.
9. Remove the wash buffer and add 70μl of secondary antibody dilution. Incubate at 4°C on a rocker at 80 rpm for 24 - 48 hours protected from light.
10. Remove the secondary antibody and add 70 μl of wash buffer and place in a rocker at 80 rpm for 10 minutes.

#### V. Nuclei staining with DAPI

11. Prepare DAPI at 1:1000 dilution from the stock in the wash buffer and incubate organoids for a minimum of 1 hour.
12. Remove the DAPI solution and add 70 μl of wash buffer and place in a rocker at 80 rpm for 30 minutes. Repeat 2 more times.

### D. Minibrain preparation for microscopy

#### I. Permeabilization of minibrain for tissue clearing

1. If the minibrains are fixed but were not processed further for immunohistochemistry, incubate organoids with wash buffer for 2 hours.

#### II. Mounting and clearing organoids for 3D confocal imaging for upright confocal microscopy

2. In the meantime, prepare the inserts for confocal imaging as shown in Figure 6. Peel the adhesive protector from the middle on one side of the imaging insert and seal the open well holes on one side using an 18 mm glass cover slip.
3. Cut the tip of a 200 μl pipette tip, transfer organoids in the chambers of the imaging insert, remove excess buffer. The chamber can hold as much as 4-5 organoids.
4. Add 20-30 μl of RapiClear™ to the well as shown in Figure 6. Make sure wells are completely filled.
5. Remove excess RapiClear™ outside the wells completely using a 20 μl pipette.
6. Peel the adhesive tape in the middle and seal the wells of the imaging insert using another 18 mm coverslips.
7. Incubate the organoids mounted with RapiClear for 24 hours at room temperature, protected from light. Note: Optionally lift the organoids using a pipette tip before sealing the top of the imaging insert. This allows easier location of the organoids during confocal imaging.

**Figure 6:**
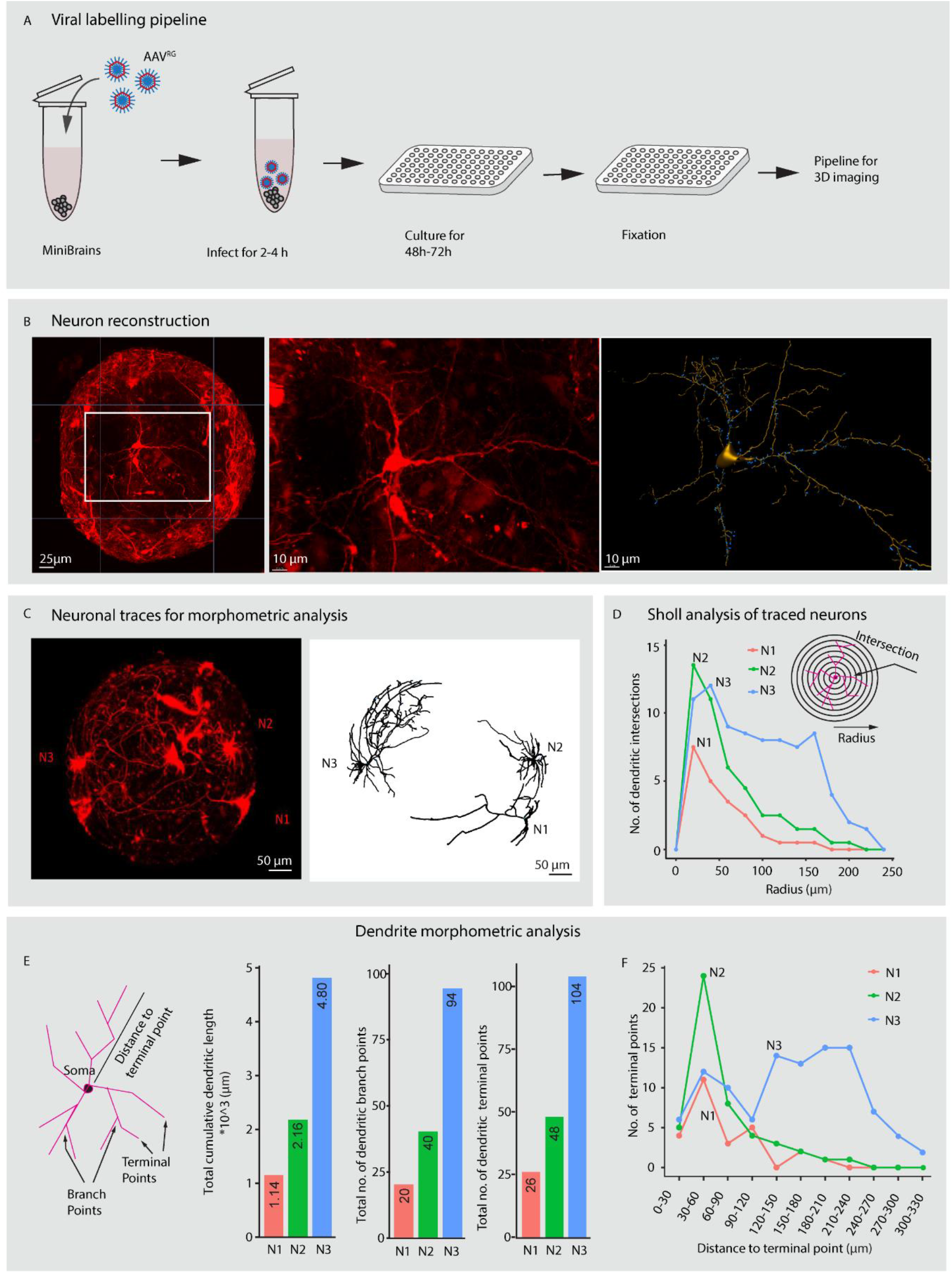
Viral labelling and neuron morphology reconstruction. **(A)** Shows a multiplexable viral infection pipeline to label projections neurons in minibrains. **(B)** Show maximum intensity projection of 3D minibrain labelled with Tdtomato by AAV-Rg infection (left), projection neuron of interest for 3d reconstruction (center) and segmented and rendered image of the neuron using filament pipeline in Imaris (Age of minibrain = 7 months). **(C)** Shows maximum intensity projection of the whole minibrain with TdTomato labelled neurons (Age of minibrain = 3 months) (left), neuron morphology traces of 3 selected neurons rendered using Imaris (right). **(D)** Sholl analysis of dendrites of the 3 selected neurons. **(E)** Shows a cartoon of dendrite morphometric features (left) and quantification of three properties. The bar graphs from left to right shows cumulative summation of all dendritic branch length, dendritic branch points and dendritic terminal in 3 traced neurons shows in panel C. **(F)** Shows distribution of dendritic terminal points at different distances from the soma across the 3 neurons traced in Panel C.

**Figure 7:**
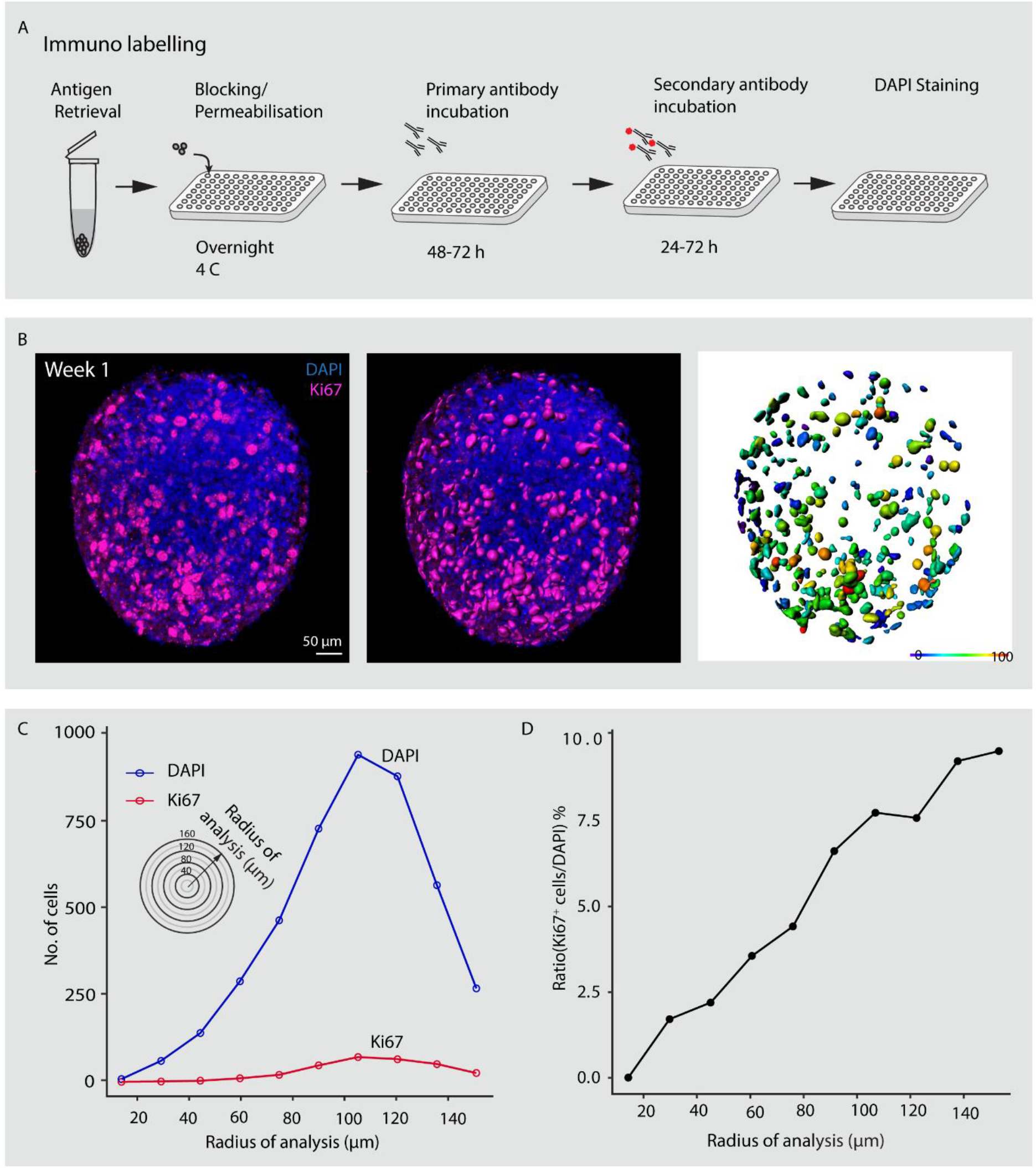
Immunohistochemical analysis of progenitors in developing minibrains. **(A)** Shows a multiplexable immunohistochemistry pipeline for minibrains. **(B)** Rendered maximum intensity projection of 3D whole minibrain image (Age of minibrain = 1 week) stained for Ki67 (progenitor marker) and DAPI (left), subsequent segmentation of Ki67 cells (center) and color-coded intensity mapped distribution of Ki67 expressing progenitors across the same minibrain 3D image (right). **(C)** Shows the number of Ki67 and DAPI stained cells across distinct radii of the sphere. **(D)** Shows the percentage of Ki67 expressing progenitors with respect to the number of DAPI stained cells, illustrating the increase in progenitor density towards the periphery.

#### III. Minibrain preparation for 3D imaging using inverted fluorescence microscopy

8. Transfer permeabilized organoids to imaging grade 96 well plate or 384 well plate.
9. Remove any remaining buffer completely and add 70 μl of RapiClear™ for 96 well plate and 40 μl of RapiClear™ for 384 well plate. Incubate in RapiClear™ overnight and proceed for imaging.

### E. Imaging organoids

#### I. Imaging in upright confocal microscopy

1. Mount 20X high resolution objective immersion lens onto the Zeiss 880 confocal microscope.
2. Prepare the mounting insert by cleaning it. Place the imaging insert as in Figure 5 and secure the insert tightly using the screws on the image insert holder.
3. Fill the mounting insert with Histodenz solution (Refractive Index =1.46)

Note: if the imaging insert is not secured using the brace, the insert will start lifting up while imaging.

4. Perform multi-tile imaging. Imaging with LSM-880 confocal microscope (Zeiss) equipped with a 20X/1 water immersion objective, with adjustable RI correction collar (RI 1.42 to 1.48). Images are collected as z-stacks using confocal or Airy scan module, with step-size of 3μm for neuron reconstructions, and 10μm for profiling the minibrain for signal.

Note: Alternatively, lightsheet microscopy technique can be used to image cluster of minibrains all at once, by using a lightsheet microscope optimized for large cleared samples (Clarity Optimized Light-sheet Microscope - COLM) (Supplementary video 2)

Note: To perform image analysis and 3D rendering on the z-stack use FIJI or Imaris software (filament tracer and surface modules). The automatic filament tracer pipeline can be applied upon background subtraction to reconstruct neuron morphology in Imaris software.

#### II. Imaging in inverted microscopy multiplexed at 96 or 384 samples

5. Choose a lens with large working distances with appropriate refractive index.
6. To ensure the organoids are close to the imaging surface, remove 30 μl of RapiClearT^M^ from each well for a 96 well plate and 30 μl of RapiClear™ from each well for a 384 well plate.
7. Proceed for imaging using required Z step size. **Troubleshooting**: If the imaging is blurry, choose appropriate RapiClear^TM^ or change the objective with the correct refractive index to match the refractive indices of both.

### Timing

#### Section A

I. Thawing frozen aliquots of NSC^hIPS^ Step 1 - Step 10: 1h 30 minutes
II. Passaging and expansion of NSChIPS Step 13 - Step −20: 1h 30 minutes
III. Week 0: Formation of spheres (3D): Step 22 - Step 26: 1 hour
IV. Week 1: Differentiation phase I: Step 27 - Step 28: 30 minutes
V. Week 2-4: Differentiation phase II: Step 29: 15 minutes for one plate
VI. Week 3-5: Shifting to maintenance phase of the minibrains: Step 30: 15 minutes for one plate
VII. ALI culture of minibrain on confetti. Step 31-37:

#### Section B

Step 1-8: 30 minutes
Step 9: 2-4 hours
Step 10-11: 48 hours

#### Section C

I. Minibrain fixation Step 1-2: 1 hour 30 minutes
II. Antigen retrieval Step 3-4: 1 hour 15 minutes
III. Permeabilization and blocking of the organoid Step 5-6: 5 hours to 18 hours
IV. Primary and secondary antibody labelling Step 7-10: 3 days
V. Nuclei staining with DAPI Step 11-12: 4 hours

#### Section D

I. Permeabilization of minibrain for tissue clearing Step 1: 1 hour 30 minutes
II. Mounting and clearing organoids for 3D confocal imaging for upright confocal microscopy Step 2-5: 20 minutes Step 6: 24 hours
III. Minibrain preparation for 3D imaging using inverted fluorescence microscopy step 7-8: 3 minutes for one well

#### Section E

I. Imaging in upright confocal microscopy Step 1-3: 20 minutes Step 4: 30 minutes to 2 hours for one organoid
II. Imaging in inverted microscopy multiplexed at 96 or 384 samples. Step 7: 1 minutes for 1 well Step 8: 10 minutes to 30 minutes per minibrain (250 μm of imaging)

## Anticipated Results

The protocol allows longitudinal maintenance of minibrains in large numbers for long periods of time. The protocol provides great control over the size of the minibrains thereby allowing a great reduction in necrosis. The viral labelling protocol is easy and quick allowing visualization of neurons as early as 24 hours after infection. Projection neurons with different morphologies are labelled through the protocol (Supplementary figure 3). Immunohistochemistry protocol is optimized to enable complete penetration of antibodies across the minibrain. The clearing of organoids is suitable for both upright and inverted microscopy allowing imaging of organoids in 3D. The protocol allows complete imaging of the minibrain from which whole neuron morphology can be reconstructed. Whole neuron morphology allows measuring complex dendrite morphometric properties.

## Limitations

Minibrain is a spheroid brain organoid, a complex ensemble of various types of excitatory neurons, inhibitory neurons and glial cells. The heterogeneity of the neurons is yet to be characterized. The size and number of organoids per well vary within and across batches of minibrains (Figure 3). The viral labelling protocol we developed is currently limited to strong ubiquitous CAG promoters. Further testing is required to check efficiency of weak promoters that would allow labelling of specific cell types. The time required for imaging the minibrains in their whole thickness using a laser scanning technique can take up to 1-2 hours when using a step size of 3 μm. This limitation can be circumvented by the use of lightsheet technology. We show that by using a clarity optimized lightsheet system (COLM), aggregates of minibrains can be quickly imaged all at once (Supplementary video 2). Localizing single organoids and vertically mounting organoids is complicated and not convenient, hence it is difficult to multiplex imaging using this kind of light sheet microscopy set up. Further protocol establishment is required for light sheet imaging of minibrains. For large screening studies, we show that clarified minibrains can be imaged using 384 well imaging culture plates (Supplementary video 5) but images can be clearly obtained only until partial depth due to the quality of the lens used that is not optimized for tissue clarified samples.

## Discussion

Minibrains are an excellent choice of *in vitro* models to study early human neuronal development, aspects of gliogenesis, neurogenesis and neuronal connectivity. The iPSCs technology allows us to generate patient specific minibrain models for various neurological disorders. Minibrains are cost-efficient, reliable and reproducible for testing drug therapeutic options, screening for toxicological effects and assaying for biocompatibility of human neural tissue. The ability to maintain the minibrains for over a year allows the researcher to monitor and follow neuronal differentiation and network maturity longitudinally over time.

In this study we present methodology for mass generation and maintenance of minibrains for large scale studies. In our methodology we adopt generation of minibrains from NSC^hiPSC^ instead of directly from iPSCs to circumvent multiple stem cell differentiation steps. We follow a slow progressive differentiation protocol that allows us to generate relatively smaller brain organoids compared to other protocols^6^. The use of breathable adhesive seals in our protocol reduces the frequency of medium changes and increased humidity maintenance, assuring better health of minibrains. ALI maintenance of minibrain allows integration of minibrains on micro-electrode array biochips and test biocompatibility of neural tissue on neuroprosthetic devices.

In this study, we present easy methodologies to study minibrains using neuronal labelling, immunohistochemistry, clearing and 3D imaging at multiplexed scales. The development of a custom multi-samples holder for the clarity confocal modules enables to image up to 9 wells, and at least 5 minibrains per well. Using immunohistochemistry and 3D imaging we show that unlike cerebral organoids, in minibrains the progenitors (expressing Ki67) are not distributed to a central core but spread throughout the minibrains (Figure 7). With our imaging pipeline, 2D imaging or partial 3D imaging of organoids can be multiplexed for up to 96 samples using inverted microscopy, investing in long distance lenses, could possibly allow whole mount organoid imaging using inverted microscopy.

It is not yet well understood how neurons shape their morphology in brain organoids where spontaneous neuronal activity based networks is the main source of input and anatomical distribution of molecules like in human developing brains is absent. Thus studying neuron morphology in brain organoids, especially the spheroid models, will allow us to study intrinsic self-organising mechanisms that guide neuron morphology across distinct neuronal subtypes. By combining viral labelling, tissue clearing and confocal imaging, we were able to produce high resolution imaging dataset from whole minibrains revealing diverse neuron morphology reconstructions. Combining neuronal markers with our protocol will allow users to reconstruct neuron morphology specific to distinct subtypes of neurons in minibrains. By using Imaris Filament tracer we were able to segment 3D labelled cells and model neuronal dendrites allowing us to map various dendrite morphometric properties of neurons ^7^. The protocol can easily be adapted for light sheet microscopy systems that have been designed for imaging small-sized biological samples and organoids, which will allow easy multiplexing and shorter time scales for imaging.

In summary the methodologies presented here will serve as a blueprint in using minibrains for large scale screening and modelling studies for the purpose of studying neuronal disorders, drug testing and chemical screening.

## Author contributions

Subashika Govindan and Adrien Roux designed and conceived the project. Subashika Govindan and Adrien Roux develop protocols and pipelines. Subashika Govindan and Samira F Osterop performed experiments. Subashika Govindan, Samira F Osterop and Laura Batti performed imaging. Subashika Govindan and Laura Batti performed image processing and analysis. Subashika Govindan wrote the manuscript. Luc Stoppini and Adrien Roux supervised the project and provided critical inputs. All authors assisted in the preparation of the manuscript.

## Acknowledgment

Laetitia Nikles and Enes Karavdic for technical and imaging support. Audrey Tissot for light sheet microscopy imaging. Loris Gomez Baisac, Jeremy Laedermann and Fabien Moreillon for engineering. Dr. Stéphane Pagès and Dr. Corinne Brana for critical input. BNF, HEPIA HES-SO, SCAHT, Wyss Center for financial support.

