## Supplementary methods and figures for "Mass Generation, Neuron Labelling and 3D Imaging of Minibrains"

### Supplementary material and methods

**Light sheet microscopy:**

Light-sheet imaging was performed using a customized version of the Clarity Optimized Light-sheet Microscope (COLM)2 at the Wyss Center Advanced Light-sheet Imaging Center, Geneva. Briefly, the sample was illuminated by one of the two digitally scanned light sheets. Emitted fluorescence was collected by a 10X XLFLUOR4X N.A. 0.6, filtered (609/54 nm, Semrock BrightLine HC) and imaged on an Orca-Flash 4.0 LT digital CMOS camera at 4 frames per seconds.

**Golgi-Cox staining of whole mount organoids**

The Ultra-rapid Golgi stain (URG) solution was prepared as described by Kassem, 2018. Minibrains were stained with URG solution in a glass vial in dark at 42°C for 24 hours. Minibrains were then washed twice in distilled water and incubated in 30% ammonium hydroxide for 20 minutes. Minibrains were then rinsed once in distilled water and were incubated in 10% sodium thiosulfate for 20 minutes. Minibrains were rinsed briefly in distilled water and incubated in DAPI (10 µg/ml in PBS with 0.1% Triton X100) for an hour. Minibrains were then washed with PBS and mounted for clearing and 3D imaging. Imaging was performed with Zeiss LSM 880 confocal microscope with reflection imaging using the 488nm wavelength, as described in Kassem et al., 2018.

**Image processing:** To perform image analysis and 3D rendering on the z-stacks, FIJI or Imaris software (filament tracer and surface modules) were used. The automatic filament tracer pipeline was applied upon background subtraction to reconstruct neuron morphology in Imaris software. Dendritic properties for Sholl analysis, number of terminal points, total dendritic lengths and branch points were retrieved from the neuronal 3D reconstructions using the Imaris software.

**Counting cells**

The position of stained cells (Ki67$^{+}$ or POU3F2$^{+}$) and DAPI cells was grouped based on their X, Y, Z coordinates in 10 concentric circles across the different radius of the sphere.- The ratio of stained cells to DAPI cells in each circle was calculated as a percentage.

**3D imaging microscopy support**

ABS-P430™ XL Model (Ivory) (Stratasys, #345-42005)

3D printer SR-30™ XL Soluble support (Stratasys, #345-42207)

The microscopy support for the sample holder was 3D printed with Uprint Stratasys 3D printer using ABS material and soluble support.

Please contact corresponding author for 3D printing source file.

**Laser cutting sample holder**

Materials:

- 2 mm Poly(methyl methacrylate) (PMMA) plate (TopAcryl Hesaglas, #VOS120000000)
- Double-sided adhesive (3M, #467MP)
- Laser cutter (Trotec, Speedy 100R)

Steps:

1. Remove the protective film on one side of the PMMA plate and apply the double-sided adhesive.
2. Repeat step 1 on the other side of the PMMA plate.
3. Place the prepared PMMA plate in the laser cutter.
4. Turn on the laser cutter.
5. Focus the laser beam on the PMMA plate and launch the software.
6. Connect the software to the laser cutter.
7. Place the laser on the desired cutting space and create an anchor point of the position in the software's workspace.
8. Import the file "adapt_Minibrain_conf_2mm_cut_and_467mp_precut.svg".
9. Drag the imported cutting procedure in the software's workspace and set the power to "PMMA VOS 2mm" from the HEPIA's directory.
10. Vectorize the cutting job.
11. Anchor the cutting job to the previously made anchor point.
12. Estimate the cutting times and launch the cutting job.
13. Allow around 1 minute to extract the gases after the end of the cutting job prior opening the hood.
14. Flip the cut piece without moving the PMMA plate.
15. Repeat steps 8 to 13 using the file "adapt_Minibrain_conf_467mp_precut_only.svg"
16. Remove the cut piece and the PMMA plate from the laser cutter.
17. Repeat steps 8 to 16 to obtain the desired number of pieces.
18. Degas the pieces using an oven at 80°C for around 72 hours.
19. The pieces are ready to be used

Note:

- Contact for "adapt_minibrain_conf_2mm_cut_and_467mp_precut.svg" and "adapt_minibrain_conf_467mp_precut_only.svg" files.

- Store the pieces in an enclosed space to prevent dust buildup
- 3D rendering of the sample holder can be visualized using creo software.

  adapt_minibrain_conf.prt.5

  Asm_adapt_minibrain_conf.asm.1

  coverslip_18mm.prt.2

  Contact for the files to print the sample holder and to visualize it.

**Supplementary Figures**

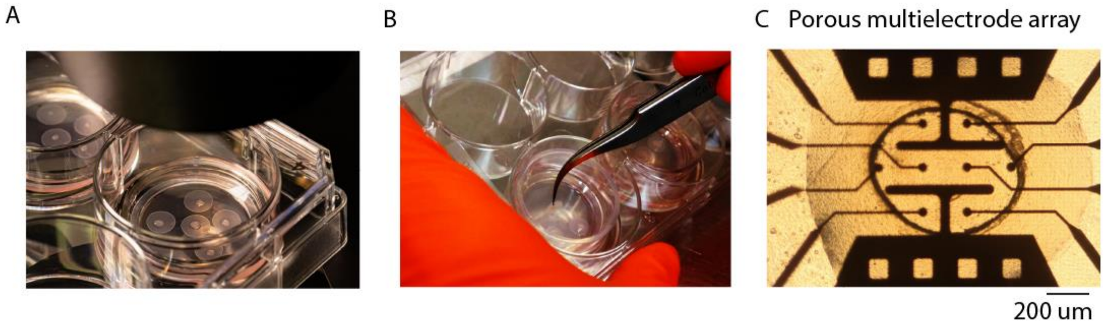


#### Supplementary figure 1:

**(A)** Shows the ALI culture of minibrains on confetti. **(B)** Shows ease of handling minibrain on confetti using forceps. **(C)** Shows minibrain on confetti integrated on biochip for electrophysiological recording.

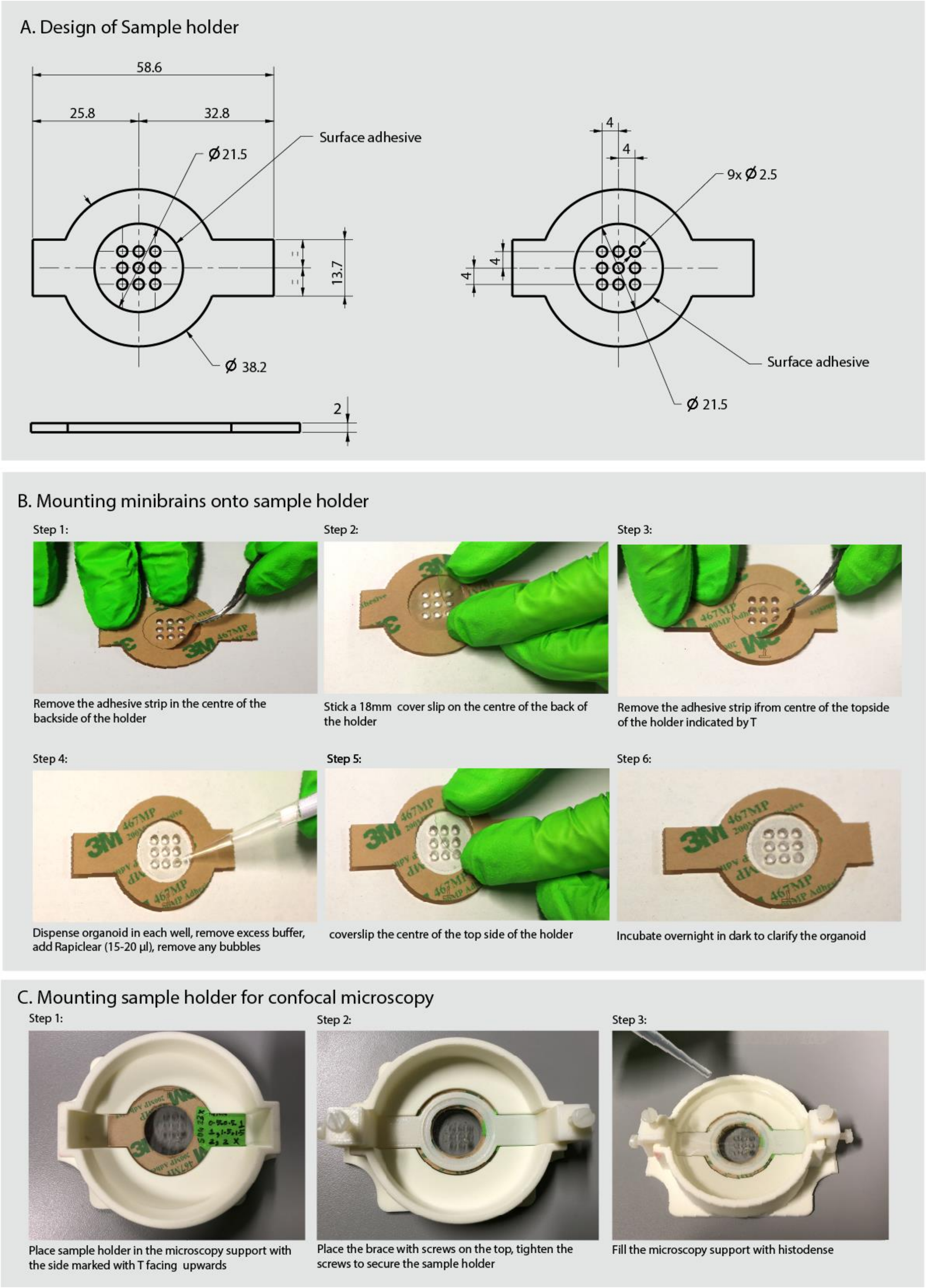


**Supplementary figure 2: Mounting organoids for 3D imaging.**

**(A)** Shows the geometry of the sample holder. **(B)** Shows mounting of minibrains on sample holder. **(C)** Shows mounting of sample holder with minibrains on microscope support for confocal imaging using the LSM 880 Zeiss microscope.

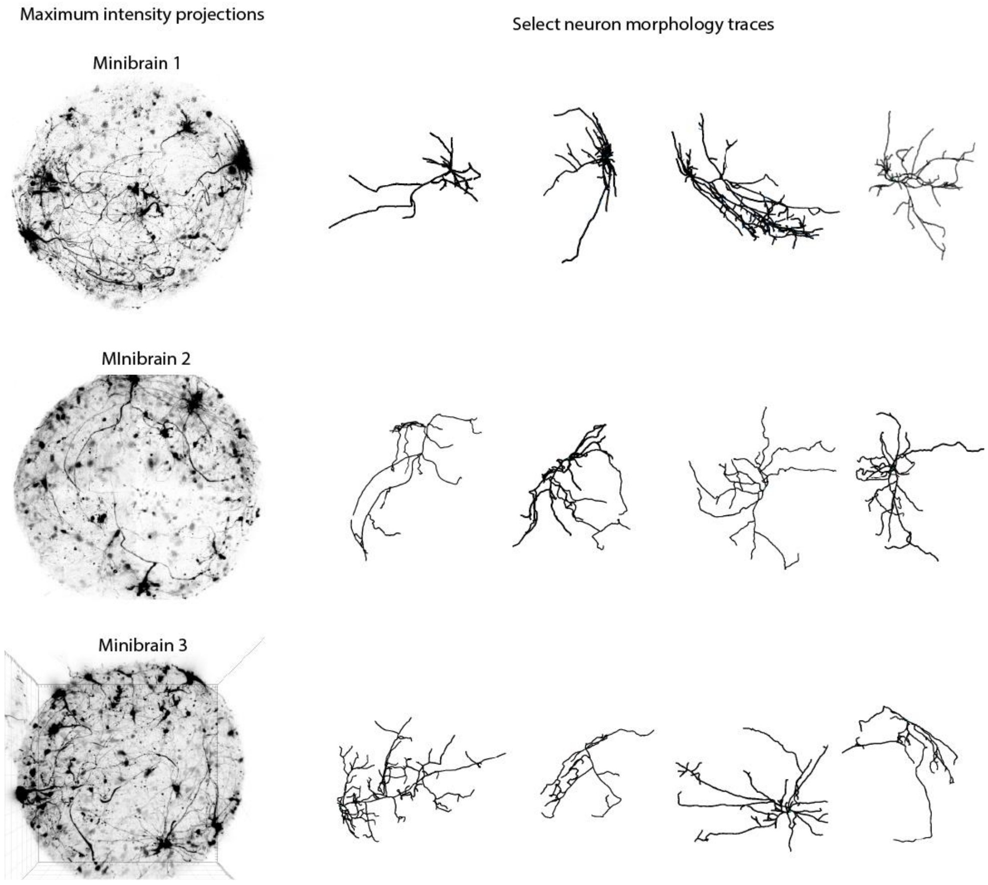


**Supplementary figure 3: Diversity in neuron morphology in minibrains**

(Left) shows maximum intensity projections 3D image of AAVrg_Tdtomato viral labelled minibrain (Age of minibrain = 3 months)-, (Right) shows select neuron morphologies from the minibrains.

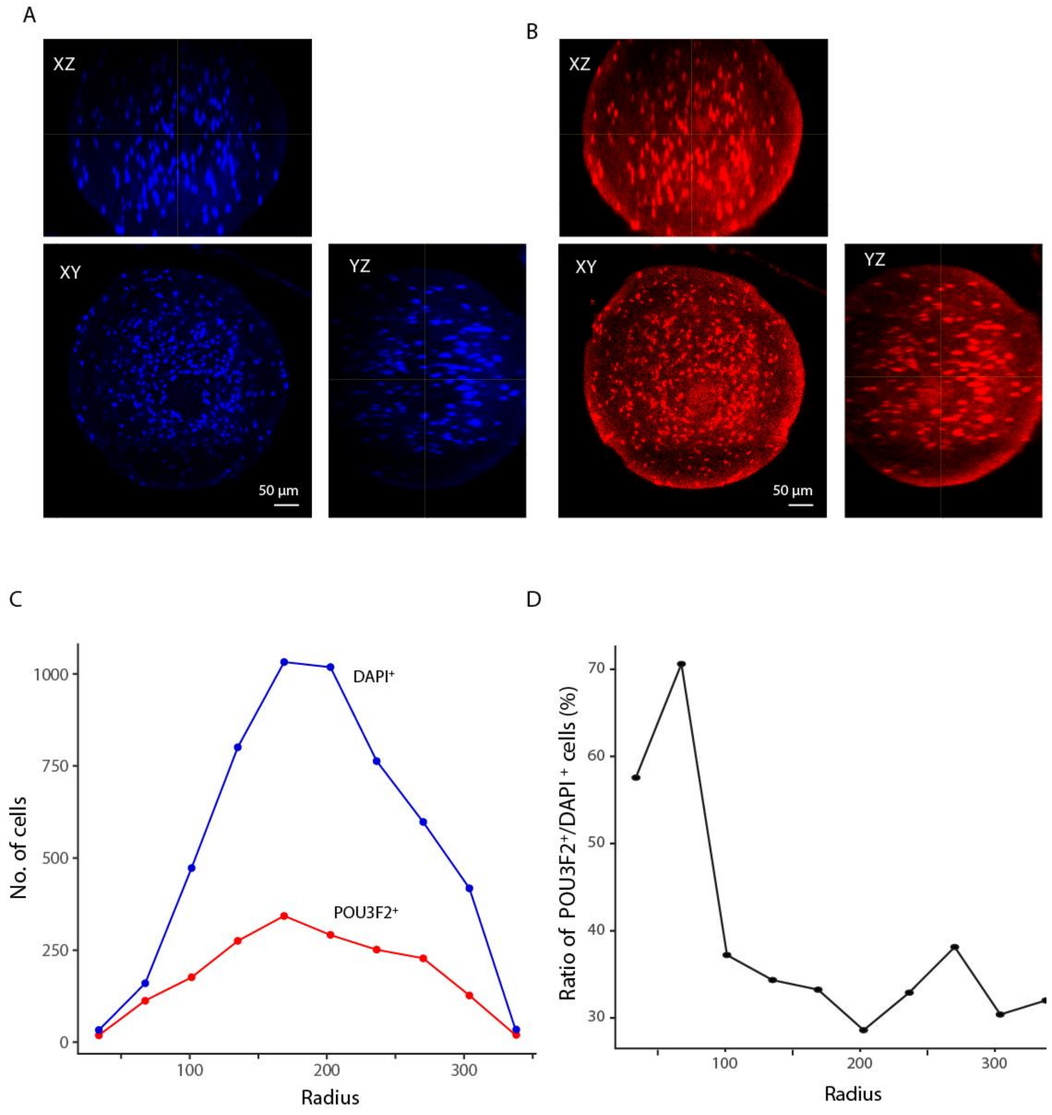


**Supplementary figure 4:**

**(A)** Orthogonal view of 3D whole minibrain image (Age of minibrain = 5 months) stained for POU3F2 (a neuronal marker) (left) and DAPI (a nuclear stain) (right) **(B)** Shows number of POU3F2 and DAPI stained cells across distinct radii of the sphere. **(C)** Shows the percentage of POU3F2 expressing neuron with respect to the number of DAPI stained cells, illustrating the dense center of the minibrain populated by the POU3F2 expressing neurons.

### Videos

**Supplementary video 1:**

Volume rendered and segmented 3D image of Golgi-Cox stained minibrains

**Supplementary video 2:**

Light sheet microscopy 3D imaging of aggregated minibrains

**Supplementary video 3:**

3D image of viral labelled neurons in 3-month-old minibrains

**Supplementary video 4:**

3D image of Ki67 staining of 1-week-old minibrain

**Supplementary video 5:**

3D image of DAPI labelled 5-month minibrain in inverted microscope with 20X air objective, scale in µm.
