## Supplementary Table 1 for "Mass Generation, Neuron Labelling and 3D Imaging of Minibrains"

### Price of medium

| EXPANSION MEDIUM | Supplier | Reference | Price /100 ml |
|---|---|---|---|
| GlutaMAX™-I Supplement | ThermoFischer | 35050038 | 1.12 CHF |
| Stempro™ NSC SFM (kit) | ThermoFischer | A1050901 | 99.00 CHF |
| | | Total | **100.12 CHF** |

| Diff1 medium | Supplier | Reference | Price /100 ml |
|---|---|---|---|
| Stempro®hESC SFM (kit) | ThermoFischer | A1000701 | 110.80 CHF |
| BDNF | ThermoFischer | PHC7074 | 157.20 CHF |
| GDNF | ThermoFischer | PHC7044 | 134.00 CHF |
| dbcAMP | Merck Sigma | D0627 | 51.20 CHF |
| 2phospho-Ascorbic Acid | Merck Sigma | 49752 | 0.01 CHF |
| | | | **453.21 CHF** |

| Diff2 medium | Supplier | Reference | Price /100 ml |
|---|---|---|---|
| Diff1 medium | | | 453.21 CHF |
| NDM medium | | | 25.04 CHF |
| | | | **478.25 CHF** |

| NDM medium | Supplier | Reference | Price /100 ml |
|---|---|---|---|
| Neurobasal ™ Plus (kit) | ThermoFischer | A3653401 | 24.76 CHF |
| GlutaMAX™-I Supplement | ThermoFischer | 35050038 | 0.28 CHF |
| | | | **25.04 CHF** |

#### Medium cost of minibrains in 6 well plate

| **Week diff** | **0** | **1** | **2** | **3** | **4** | **5** | **6** | **7** | **8** | | 52 |
|---|---|---|---|---|---|---|---|---|---|---|---|
| **Medium** | Prolif | Diff-1 | 50-50 Diff-2 | | | NDM | | | | | |
| **Price/Well/week** | 3.00 CHF | 11.33 CHF | 5.98 CHF | 5.98 CHF | 5.98 CHF | 0.63 CHF | 0.63 CHF | 0.63 CHF | 0.63 CHF | | |
| **Total/well** | | | | | | | | | **34.77 CHF** | | **62.32 CHF** |
| **Total for 10 plates** | | | | | | | | | ########### | | ########### |
